# Non-neutral evolution of H3.3-encoding genes occurs without alterations in protein sequence

**DOI:** 10.1101/422311

**Authors:** Brejnev M. Muhire, Matthew A. Booker, Michael Y. Tolstorukov

**Affiliations:** Department of Molecular Biology, Massachusetts General Hospital and Harvard Medical School, Boston, MA 02114

## Abstract

Histone H3.3 is a developmentally essential variant encoded by two independent genes in human (*H3F3A* and *H3F3B*). While this two-gene arrangement is evolutionarily conserved, its origins and function remain unknown. Phylogenetics, synteny and gene structure analyses of the H3.3 genes from 32 metazoan genomes indicate independent evolutionary paths for *H3F3A* and *H3F3B*. While *H3F3B* bears similarities with H3.3 genes in distant organisms and with canonical H3 genes, *H3F3A* is sarcopterygian-specific and evolves under strong purifying selection. Additionally, *H3F3B* codon-usage preferences resemble those of broadly expressed genes and ‘cell differentiation-induced’ genes, while codon-usage of *H3F3A* resembles that of ‘cell proliferation-induced’ genes. We infer that *H3F3B* is more similar to the ancestral H3.3 gene and likely evolutionarily adapted for broad expression pattern in diverse cellular programs, while H3F3A adapted for a subset of gene expression programs. Thus, the arrangement of two independent H3.3 genes facilitates fine-tuning of H3.3 expression across cellular programs.

## Introduction

In eukaryotic cells genomic DNA is packaged into chromatin, which plays a dual role of genome compaction and regulation [1]. Basic repeating units of chromatin, called nucleosomes, comprise 147bp of DNA wrapped around a core that is formed by histone proteins of four types (H2A, H2B, H3, and H4), which are conserved from yeast to human [2,3]. The histones fall into two major types: replication-dependent (RD) canonical histones and replication-independent (RI) non-canonical variants. The RI histone variants have diverse biological roles and are part of epigenetic regulation of genome function [4–6]. Unlike the canonical histones that are encoded by co-regulated gene clusters (histone loci) [3], RI variants are encoded by individual genes that are regulated similarly to other protein coding genes.

One of the most studied histone variants is H3.3, which replaces canonical histone H3 and functionally can be associated with both gene activation [7,8] and silencing [9–11]. H3.3 variant is expressed and deposited throughout the cell-cycle independent of DNA replication [12–14]. In human genome H3.3 can be transcribed from either of two independent genes (*H3F3A* and *H3F3B*), which are located at different chromosomes, 1 and 17 respectively. These genes differ at the nucleotide level both within introns and exons, even though both of them encode exactly the same amino-acid sequence. Presence of multiple independent genes encoding H3.3 is also conserved in other organisms, including distant species such as fruit fly [15]. Moreover, despite absolute conservation at the protein level, the mutational profiles of *H3F3A* and *H3F3B* genes in human cancers differ substantially. For instance, mutation K27M was reported in only in *H3F3A* in brainstem gliomas [16], while mutation K36M is predominantly observed in *H3F3B* in bone cancers, such as chondroblastoma [17,18]. The regulatory genomic elements associated with these genes are also distinct, and the over-expression of *H3F3A* but not *H3F3B* is implicated in lung cancer through aberrant H3.3 deposition [19]. Taken together, these observations indicate that while *H3F3A* and *H3F3B* encode the same protein product, they are under different regulatory mechanisms and play distinct roles.

Evolution of H3.3 encoding genes was analyzed in *Drosophila* species [20], however, on a larger scale, the biological function and evolutionary history of such two-gene organization remains unclear, despite its biomedical significance [21,22]. To approach these questions, we compared the sequences and genomic arrangements of the H3.3 genes from 32 metazoan genomes. Using phylogenetics, sequence identity, gene structure and synteny analyses we infer that *H3F3A* is sarcopterygian-specific (tetrapod and lobe-finned fish) gene, while *H3F3B* is of more ancient origin. Furthermore, analysis of codon-usage preferences in each of the H3.3 genes revealed that *H3F3B* is evolutionarily adapted for broad expression patterns across diverse cellular programs, including cell differentiation, while *H3F3A* is more fine-tuned for a specific transcriptional program associated with cell proliferation. This observation of coding sequence optimization for distinct transcriptional programs provides insight into why both *H3F3A* and *H3F3B* have been maintained in course of evolution, even though they produce identical proteins.

## Results

### Phylogenetic analyses of H3.3-encoding genes in metazoa

We identified the H3.3 coding sequences from the genomes of 32 metazoa organisms, primarily vertebrates, and used them in our analysis. We observed that two ‘independent’ genes (i.e. located in different genomic loci and controlled by distinct, non-overlapping promoters) encode histone H3.3 in all analyzed organisms except for actinopterygii (ray-finned fish lineage) and coelacanth where H3.3 is encoded by three or five genes (Table S1). The high number of H3.3 genes in actinopterygians most likely resulted from whole genome duplication events [23–26] and partial chromosome duplication events [27–29] that occurred in this lineage during evolution. With this exception, the arrangement of two H3.3 genes is widespread among vertebrates and is observed even in more distant metazoa such as flies, nematodes, and some plants [30]. Remarkably, the encoded protein sequence is identical in all vertebrates and *Drosophila melanogaster* (Fig S1). The existence of two independent genes that encode an identical protein allows us to focus on analysis of the evolutionary pressure acting on these genes at the nucleotide rather protein level.

Next, we analyzed phylogenetic relationship of the H3.3 genes in metazoa. The coding sequences of these genes form several distinct groups in the phylogenetic tree, including two major groups (clades 1 and 3), one minor group (clade 2) and outgroups of lamprey and fly H3.3 genes (Fig. 1A). Clade 1 (shown in brown) consists exclusively of sarcopterygian *H3F3A* genes (the lobe-finned fish lineage, including all tetrapods and coelacanth). Clade 3 comprises all sarcopterygian *H3F3B* genes (blue) along with the majority of actinopterygian H3.3 genes (gray) and the third coelacanth H3.3 gene. We note that this clade also includes a ‘hominid-specific’ gene H3F3C (green), which emerged as a recent retro-transposition of *H3F3B* [31]. *H3F3C* encodes another replacement histone from H3 family, H3.5, that differs from the histone H3.3 by several amino-acids, and it was included in this analysis for further comparison. The confident assignment of *H3F3C* to clade 3 that contains H3F3B genes (branch support=1), highlights that the distinction between the coding sequences (CDS) of the genes forming clades 1 (*H3F3A*) and 3 (*H3F3B*) is substantial and evolutionary stable even though these genes encode the same protein H3.3 (no amino-acid difference). Finally, clade 2 contains remaining actinopterygian H3.3 genes that cluster neither with sarcopterygian *H3F3A* nor with sarcopterygian *H3F3B*. This analysis gives the first evidence that, compared to sarcopterygian *H3F3A*, sarcopterygian *H3F3B* is likely more evolutionarily related to actinopterygian H3.3 genes.

**Figure 1.**
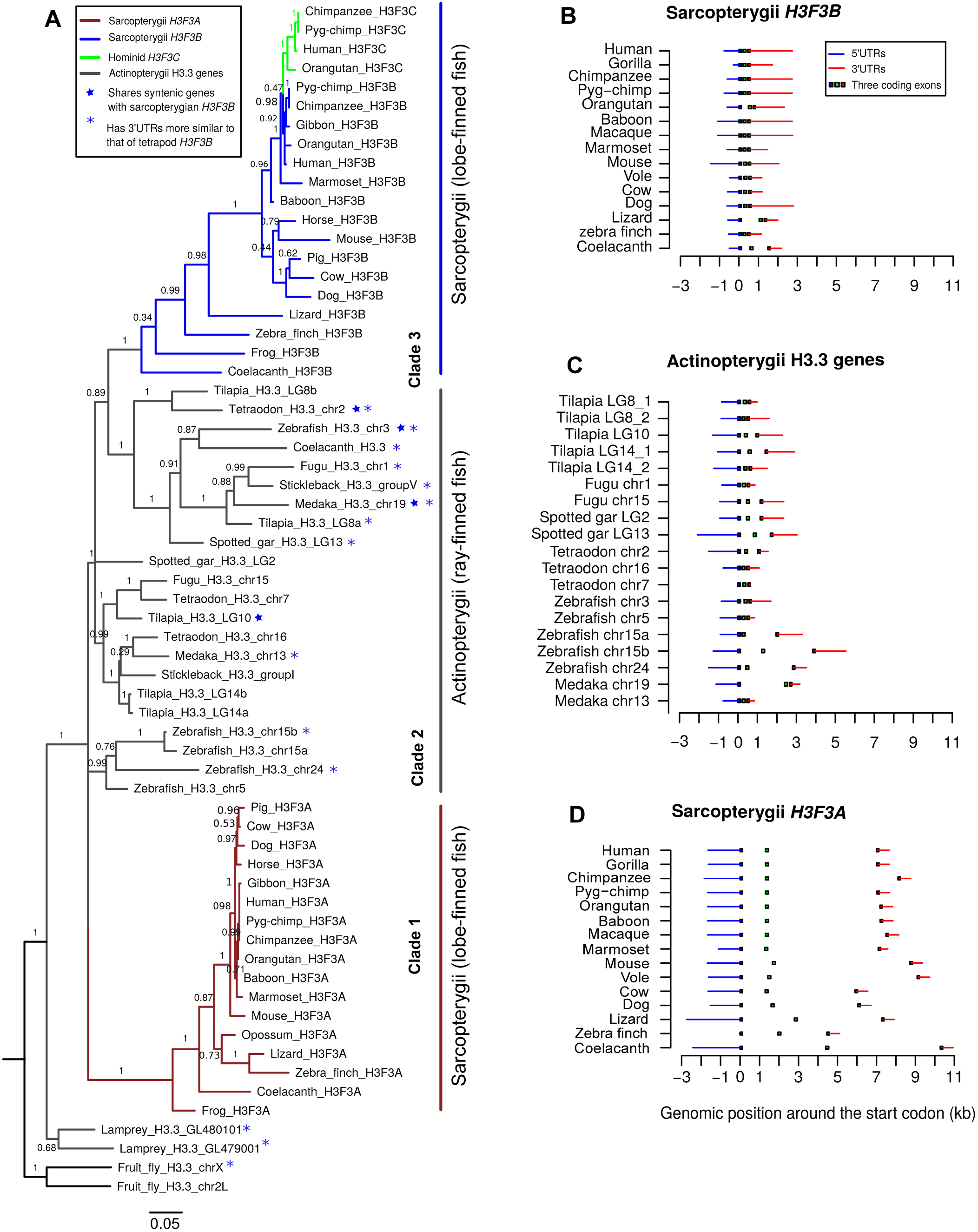
Phylogenetic analyses H3.3-encoding genes. **A.** Maximum likelihood tree illustrating the evolution of H3.3 genes in vertebrates. Three clades were distinguished. Clade 1 comprises sarcopterygian *H3F3A* genes (brown); Clade 3 comprises sarcopterygii *H3F3B* (blue) which cluster together with actinopterygian H3.3 (gray). Clade 2 consists of other actinopterygian H3.3 genes that cluster with neither clade 1 nor clade 3. Numbers along tree branches represent approximate log-likelihood ratio test values for branch support. Blue stars mark non-tetrapod genes with syntenic relation to tetrapod *H3F3B*, and blue asterisks mark non-tetrapod genes whose 3’UTRs are more similar to 3’UTRs of tetrapod *H3F3B* than those of tetrapod *H3F3A*. **B, C, D.** Intron-exon structure of sarcopterygian *H3F3B*, actinopterygian H3.3 genes and sarcopterygian *H3F3A*. All genes are drawn from 5’ to 3’ and are aligned at the start codon, position 0. The blue and red lines represent the 5’ UTRs and 3’ UTRs respectively, and the squares in the middle represent the locations of protein-coding exons.

The observed relation between *H3F3B* and actinopterygian genes was confirmed by comparison of the intron-exon structure of all H3.3-encoding genes throughout the species. In sarcopterygian genomes *H3F3B* is generally shorter, spanning ∼2-4kb with a total length of introns ∼0.16-1kb (Fig. 1B). *H3F3B* structure is similar to that of actinopterygian H3.3 (gene length is approximately ∼2-6kb and total intron length is ∼0.16-4kb; Fig. 1C). The *H3F3A* gene structure is noticeably different, with gene length spanning ∼9-13kb and total intron length being ∼4.5-10kb (Fig. 1D). Thus, the intron-exon structure of sarcopterygian *H3F3B*, and not sarcopterygian *H3F3A* is more similar to the actinopterygian H3.3 genes and H3.3 genes in more distant vertebrates actinopterygians, lamprey, fly and worm, consistent with our previous observations.

To further support these results, we carried out synteny analysis to determine whether genes around *H3F3A* or *H3F3B* are evolutionary conserved in non-tetrapod organisms. We first used Genomicus 80.01, a web-based synteny visualization tool that uses Ensembl database comparative genomic data [32]. Comparison between human and actinopterygii shows no syntenic genes conserved around human *H3F3A* and H3.3 genes in actinopterygian species (Fig 2A), but at least six syntenic genes can be identified around human *H3F3B* and H3.3 genes in four actinopterygian species (fugu, platyfish, spotted gar, and tetraodon) (marked with a blue star, Fig 2B).

**Figure 2:**
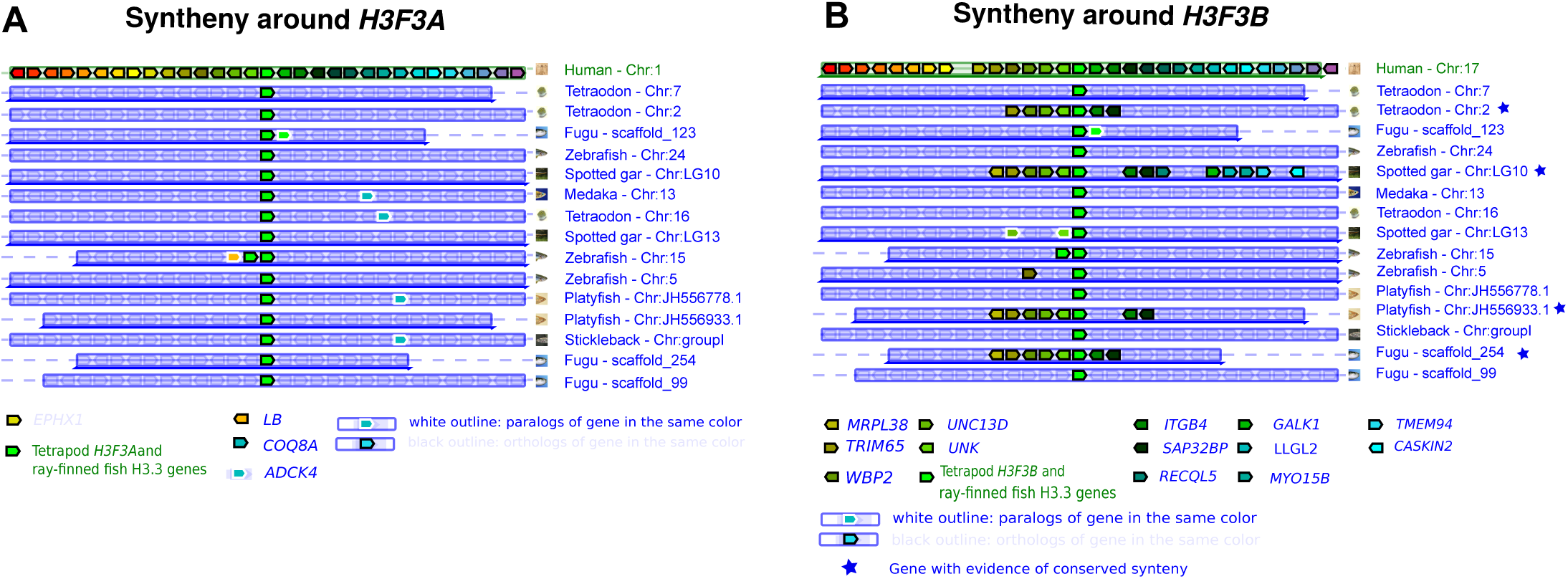
Synteny around H3F3A and H3F3B genes **A, B.** Synteny conservation analysis around human H3F3A (A) and H3F3B (B) genes performed using selected actinopterygian genomes. Human *H3F3A* and *H3F3B* and actinopterygian H3.3 are placed at the center of each plot (green block). A black outline represents an ortholog of a gene in the same color, while a white outline represents a paralog of gene in the same color. A blue star indicates actinopterygian organisms in which syntenic genes around the H3.3 gene are also conserved around the human H3.3 gene.

We extend this analysis to all tetrapods and distant metazoa (lamprey, fly and worm), by implementing a flexible synteny detection method allowing the user to quantitatively measure the degree of gene conservation around loci of interest in two genomes (see Methods). Specifically, we compared 30 genes upstream and downstream of each of the H3.3 genes and the degree of gene conservation was determined by sequence identities computed independently for both coding sequences and translated amino-acid sequences. While we found clear evidence of synteny conservation around both H3.3 genes in tetrapods, it was consistently higher around *H3F3A* than *H3F3B*. For instance, the ratios of syntenic genes around *H3F3A* to those around *H3F3B* were 25/17, 12/6, 12/6 for the human-mouse, human-lizard and human-zebra finch comparisons respectively (Fig S2A). At the same time, we found no synteny conservation around tetrapod *H3F3A* and actinopterygians H3.3. In contrast, for *H3F3B* we found the same six genes conserved between tetrapods and one of the tetraodon H3.3 genes, which were detected by Genomicus, and a weak conservation of these genes in zebrafish and medaka (marked with a blue star Fig S2B and Fig 1A).

From these observations, we conclude that orthologs of mammalian *H3F3A* and *H3F3B* are present in the coelacanth genome (i.e. throughout the sarcopterygian lineage). Sarcopterygian *H3F3B* is evolutionarily related to many actinopterygian H3.3 genes while sarcopterygian *H3F3A* seems to have no counterpart in actinopterygian lineage (Fig 1A). We infer that the sarcopterygian-specific *H3F3A* clade with a long and well-supported branch (branch support=1, Fig 1A) is consistent with one of the following scenarios: (i) the counterpart of *H3F3A* was lost in actinopterygian lineage soon after actinopterygian-sarcopterygian split, or (ii) since the actinopterygian/sarcopterygian split either an existing or a newly emerged H3.3 gene underwent rapid evolution towards the current *H3F3A* form. We aimed to distinguish these possibilities by the analysis described below.

### Comparison of H3.3 genes between sarcopterygians and distant metazoa

One can expect that if *H3F3A* were lost in actinopterygians, both *H3F3A* and *H3F3B* would exhibit roughly equal similarity to H3.3 genes in more distant metazoa. Thus, to resolve the scenarios described above we directly compared the similarity of sarcopterygian *H3F3A* and *H3F3B* to the H3.3 genes of actinopterygians and distant organisms (lamprey and fly) (Fig. 3). We also included in this analysis genes encoding the RD canonical histones H3.1 and H3.2 because these genes emerged from ancient gene duplication event that resulted in a separation of replication-dependent and replication-independent histones [33]. As sarcopterygian genes in this analysis, we used coelacanth *H3F3A* and *H3F3B*. Coelacanth can be expected to show more similarity with distant organisms than other sarcopterygians, in part because its protein-coding genes evolved twice as slow as such genes in tetrapods [34], which makes it especially suitable for this comparison.

**Figure 3.**
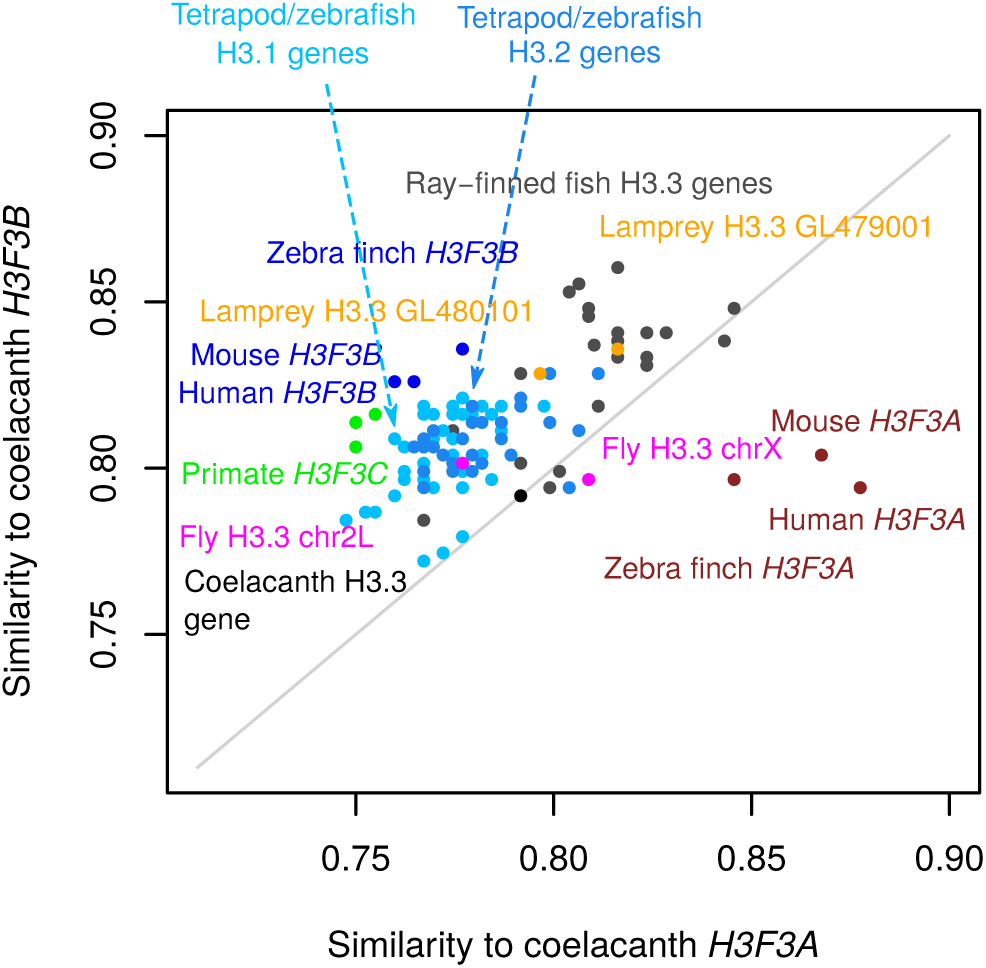
Comparison of coelacanth H3.3 genes to related genes in sarcopterygian and non-sarcopterygian lineages. Sequence similarity was estimated for the CDS of coelacanth H3.3 genes (*H3F3A,* x-axis and *H3F3B,* y-axis) and CDS of H3.3 genes from other sarcopterygian and more distant organisms (actinopterygian, lamprey, fly). Additionally, CDS of tetrapod and zebrafish H3.1 and H3.2 genes were included in this analysis. Each point represents a gene and the organism name is written in the matching color. The sequence similarity represents percentage of the identical nucleotides in the sequence.

This analysis revealed that most of the actinopterygian H3.3 genes and RD H3.1 and H3.2-encoding gene of bony vertebrates (tetrapods and zebrafish) are more similar to sarcopterygian *H3F3B* than to *H3F3A* (Fig. 3). This trend further extends to both lamprey H3.3 genes and one fly H3.3 (chr2L) gene. In addition, *H3F3C* is also more similar to coelacanth *H3F3B* than *H3F3A* as expected. Overall, only tetrapod *H3F3A* genes can be confidently ‘assigned’ to coelacanth H3F3A. As a control, we have repeated this analysis using tetrapods (human, mouse and zebra finch) *H3F3A* and *H3F3B* genes instead of coelacanth genes and observed similar trends (Fig. S3). Overall, these results reveal that in comparison to *H3F3A*, sarcopterygian *H3F3B* is more similar to the H3.3 genes in distant metazoa and to RD H3 genes, pointing to a possibility that *H3F3B* is more similar to the ancestral form of the H3.3 gene.

Additional evidence supporting the hypothesis formulated above comes from the comparison of the 3’ untranslated regions (3’UTRs) of the H3.3 genes (Fig S4). UTRs are among the most conserved non-coding sequences in eukaryotes [35,36], and the 3’UTRs of H3.3 genes are similarly evolutionarily conserved (∼60-80%) among tetrapods and actinopterygians. We validated this approach by confirming that it produces results consistent with the phylogenetic analysis of the coding H3.3 sequences when applied to genes from clades 1 and 3 (Fig. 1A), containing sarcopterygian H3.3 genes. When we applied this approach to genes from other clades, we observed that in every analyzed non-sarcopterygian organism (actinopterygian species, lamprey, fly and worm), at least one H3.3 gene has higher similarity of its 3’UTRs to that of tetrapod *H3F3B* (∼75% identity) compared to tetrapod *H3F3A* (∼60% identity) (Fig. S4AB). These organisms are marked with blue asterisks in Fig 1A. There were no instances of a non-tetrapod H3.3 3’UTR being more similar to the 3’UTR of tetrapod *H3F3A*.

Collectively, our results indicate that gene *H3F3A* is sarcopterygii-specific, while gene *H3F3B* is evolutionary related to actinopterygian H3.3 genes as well as to the H3.3 genes in more distant metazoans. Furthermore, our results suggest that *H3F3B* is more directly related to the ancestral form of the H3.3 gene. We find that the possibility of a lineage-specific loss of *H3F3A* in the actinopterygians is less plausible than the hypothesis of an existing or newly emerged H3.3 gene copy that underwent rapid evolution to become *H3F3A* in sarcopterygian lineage.

### Distinct selection pressures within tetrapod *H3F3A* and *H3F3B* CDS

The conservation of the arrangement of two distinct genes encoding the same protein suggests functional significance. To investigate how potential functional differences between these two genes may be reflected in their genomic sequences, we measured selection pressure operating at the nucleotide level in *H3F3A* and *H3F3B*. Due to lack of variation among H3.3 protein sequences in analyzed organisms, the methods based on non-synonymous and synonymous substitution rates often used for detection of natural selection [37–39] are not suitable. Instead, we investigate purifying selection operating on *H3F3A* and *H3F3B* genes based on the degree of conservation of coding nucleotide-sequence in tetrapod organisms.

We started with calculating pairwise genetic distances between the tetrapod H3.3 genes, defined here as the numbers of the observed nucleotide substitutions divided by the CDS length (i.e. the “nucleotide substitution scores”). As a control, we also included in this analysis the *H2AFZ* gene, which encodes the conserved replacement histone H2A.Z. Overall, we observed that while *H3F3B* is not conserved significantly stronger than *H2AFZ* (P = 0.244, Mann-Whitney’s test), *H3F3A* is under a stronger selection pressure as compared to both *H3F3B* and *H2AFZ* (P = 2*10^−7^, P = 3*10^−6^ respectively, Fig. 4A). Also, the distributions of the nucleotide substitution scores are bimodal for all three genes, revealing that they are particularly conserved within two distinct groups: (i) mammals and (ii) reptiles, birds and amphibians (Fig. 4A). This trend is especially pronounced for *H3F3A*, and we further confirmed a stronger conservation of this gene within each individual group of organisms (Fig. S5A-B).

**Figure 4.**
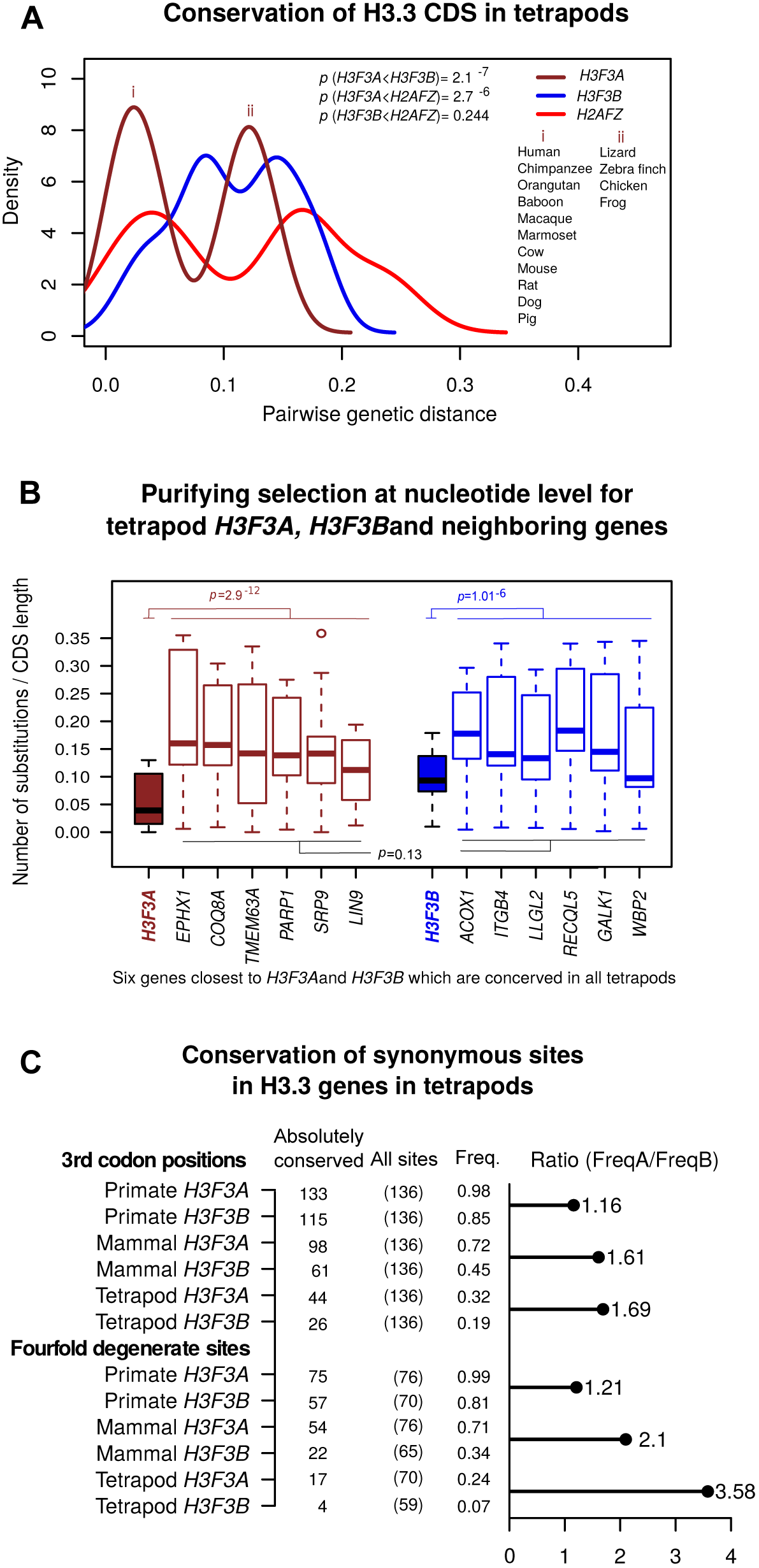
Conservation of coding sequences in tetrapod histone variant genes. **A.** Pairwise nucleotide substitution scores (genetic distances) computed for two H3.3 genes (*H3F3A, brown* and *H3F3B,* blue), and *H2AFZ* gene (red) which was included in this analysis for comparison. The analysis was performed for tetrapod genomes. Distribution shifting to the left (smaller genetic distances) indicates higher conservation of a corresponding gene. (i) marks the peak of the bimodal distribution corresponding to pairwise scores involving mammalian organisms, while (ii) shows the distribution corresponding to pairwise scores involving exclusively *H3F3A* genes in non-mammals. **B.** Pairwise nucleotide substitution scores for *H3F3A* in tetrapod genomes (box plot filled in brown), *H3F3B* in tetrapod genome (filled in blue), and their neighboring genes (brown and blue borders respectively). Both *H3F3A* and *H3F3B* are significantly highly conserved relative to their surrounding genes (Wilcox sum rank test P = 2.9^−12^ and P = 1.01^−6^ respectively). No significant difference in conservation level between genes around *H3F3A* and those around *H3F3B* (P = 0.13). **C.** Absolute nucleotide conservation in CDS of the H3.3 genes in tetrapod lineages. Top panel: all 3^rd^ codon position; bottom panel: the fourfold degenerate sites (i.e. sites where any possible nucleotide substitution is synonymous). Columns show the number of absolutely conserved sites for a given group of organisms, the total number of 3^rd^ codon positions or fourfold degenerate sites, and the corresponding frequencies of absolutely conserved sites. The horizontal bar represents the H3F3A/H3F3B over-representation of absolutely conserved sites.

To rule out that the difference in sequence conservation of H3.3-encoding genes is determined by the conservation of entire loci encompassing *H3F3A* or *H3F3B*, rather than these genes themselves, we extended the analysis described above to six genes around each of the H3.3-encoding genes. We found no significant difference in conservation level between genes around *H3F3A* and those around *H3F3B* (Fig. 4B).

At the same time, both *H3F3A* and *H3F3B* are significantly more conserved than the neighboring genes (P = 3*10^−12^ and P = 10^−6^ respectively), with *H3F3A* exhibiting highest level of conservation among the analyzed genes. This indicates that tetrapod *H3F3A* evolves under stronger purifying selection at nucleotide level than *H3F3B*, *H2AFZ* or neighboring genes.

Given that the H3.3 genes encode the same amino-acid sequence, not surprisingly most substitutions were observed in the 3^rd^ position of the codon. Interestingly, we found that sarcopterygian *H3F3B* have generally higher GC-content at 3^rd^ codon position (GC3) as compared to sarcopterygian *H3F3A* (Fig. S6). The high GC3 in *H3F3B* genes mirrors actinopterygian H3.3 and RI H3.1/H3.2-encoding genes while the *H2AFZ* genes have low GC3 that close to that of *H3F3A* (Fig S6). Thus, based on this metric *H3F3B* is more similar to ancestral H3.3 and RI H3 histone genes, hence these results are in agreement with our previous phylogenetic analyses.

To refine this analysis further, we compared the degree of nucleotide conservation at wobble positions (i.e. 3^rd^ codon positions where synonymous nucleotide substitutions are commonly detected) between *H3F3A* and *H3F3B* gene alignments made of (i) all tetrapods, (ii) mammals, and (iii) primates (Fig. 4C). We also separately considered a special case of wobble positions, so-called ‘fourfold degenerate’ sites, i.e. 3^rd^ codon positions at which all possible nucleotide substitutions can occur without changing the encoded amino-acid; hence such fourfold degenerate sites are under no selection pressure for amino-acid maintenance. A wobble position was considered “absolutely conserved” if the nucleotide at that site is conserved in the whole alignment (i.e. in all organisms).

In all groups, we consistently observed that there are more absolutely conserved 3^rd^ codon positions in *H3F3A* than *H3F3B* in all analyzed groups of species (Fig. 4C). This trend is most pronounced for the fourfold degenerate sites (cf. horizontal bars in Fig. 4C). In addition, such an over-representation is more pronounced for groups containing evolutionary distant organisms e.g. FreqA/FreqB ratio for fourfold degenerate sites is 1.21, 2.10, 3.58 for primates, mammals, and tetrapods respectively. This observation suggests that stronger selection on synonymous sites in *H3F3A* than *H3F3B* is a stable phenomenon, deeply rooted in the tetrapod lineage.

These findings revealed that there is a layer of selection pressure against nucleotide substitutions operating on both *H3F3A* and *H3F3B* CDSs, driven not by the maintenance of amino-acid sequence but maintenance of specific codons. Thus, our results suggest that codon usage is under selection pressure among H3.3 genes. While this selection pressure is stronger in *H3F3A* than in *H3F3B*, we infer that both genes have evolutionary adapted for distinct codon usage preferences, and we investigate this phenomenon in more detail below.

### Differences in codon usage between H3.3 encoding genes

The expression and abundance of transfer RNA (tRNA) vary substantially in human cell types [40]. This variation correlates with codon usage preferences and plays a role in translational control [41–43]. Furthermore, codon usage may differ between genes specialized in different cellular processes such as cell proliferation and cell differentiation [41]. Thus, an analysis of the codon usage in H3.3 genes can provide information on their functional specialization among cellular gene expression programs.

To this end, we estimated the correlation between codon usage frequencies in each of the H3.3 genes and the genome-wide codon usage frequencies from each tetrapod genome. Similar to a previously published study [41], we define these codon usage frequencies (hereby referred to as “amino-acid specific codon frequencies”) so that they represent the probability that a codon is used when the amino-acid encoded by this codon appears in the protein product sequence (see Methods). Since different genes are expressed in different cell types, we expect that the codon usage frequencies computed for the entire genome (‘genome-wide codon usage frequencies’) would correlate strongly with the codon usage frequencies of the genes showing broad expression patterns. In line with this hypothesis, codon usage frequencies in a set of human genes specifically selected for their ubiquitous expression in multiple cell types [44] correlated with genome-wide frequencies with the Pearson’s correlation coefficient equal about 0.695 (Fig. 5A). Application of this approach to the H3.3 genes revealed that the correlation estimated for the human *H3F3B* gene (r=0.69) is close to benchmark value observed for the ubiquitously expressed genes (UEG), while the correlation for the *H3F3A* gene is considerably lower (r=0.54). Furthermore, all tetrapod *H3F3B* genes, actinopterygian H3.3 genes, and RC H3.1/H3.2 genes (the latter are expressed in all dividing cells) show higher correlation with genome-wide frequencies than either *H3F3A* or *H2AFZ* genes do (Fig 5A). We confirmed that similar results are observed when codon usage is defined directly as the frequency of every codon in a gene, without accounting for amino-acid abundance in the product (“codon frequencies” in Fig. S7A). Based on these findings, we conclude that, as compared to *H3F3A*, *H3F3B* is evolutionarily more optimized for a broad expression pattern.

**Figure 5.**
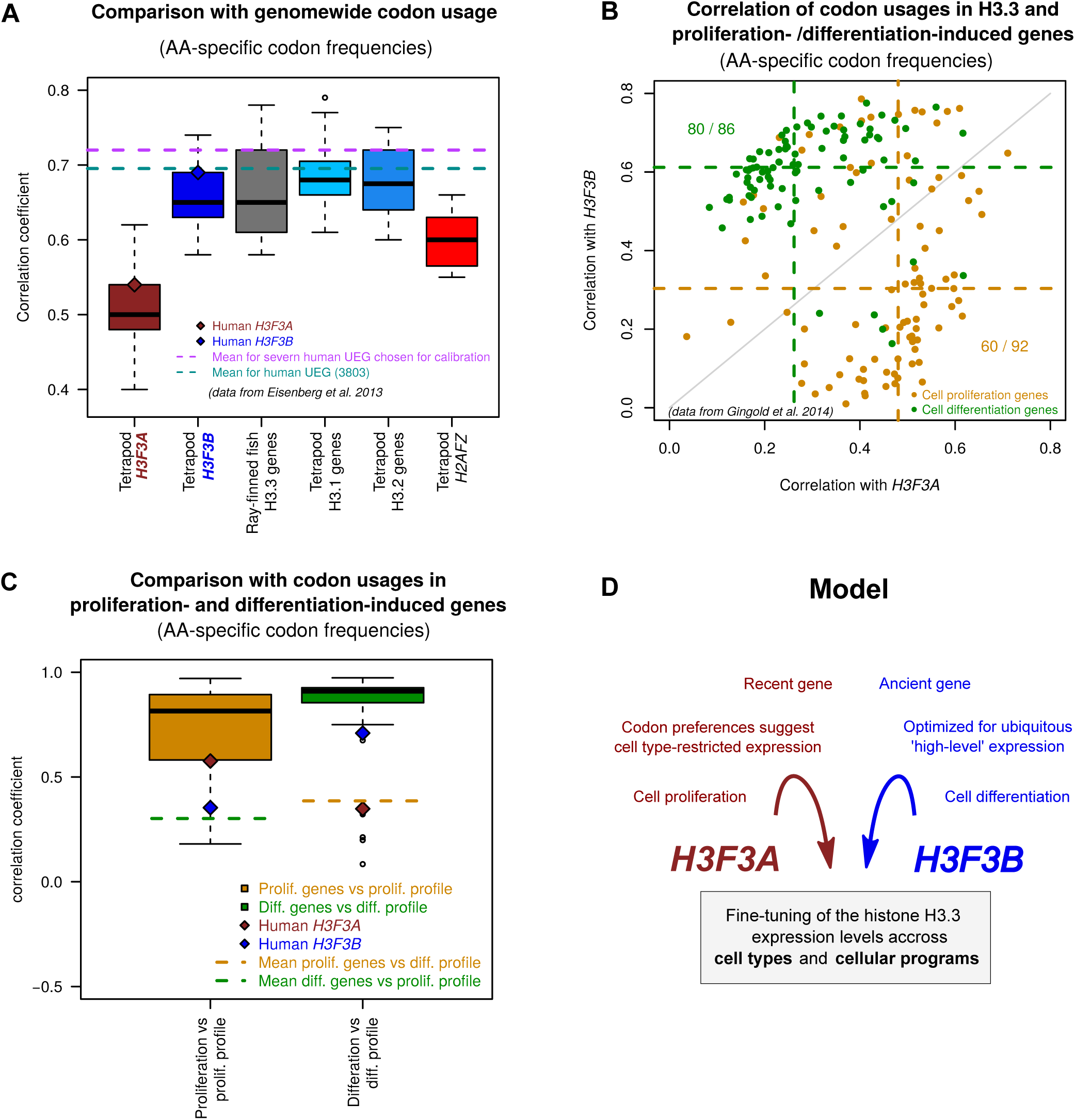
Distinct codon usage preferences in the H3.3 genes (based on ‘amino-acid specific codon frequencies’) **A.** Correlation between codon usage in the genes specified at x-axis and the genome-wide codon usage. The box plots represent the lineage distributions of the correlation coefficients calculated for the ‘amino-acid specific codon frequencies’ of a corresponding gene with those estimated genome-wide (e.g. all tetrapod H3F3A genes vs. genome-wide frequencies). The brown and blue diamonds provide reference for human *H3F3A* and *H3F3B* respectively. The pink dashed line represents average correlation computed for human ubiquitously expressed genes (UEG) [44]. **B.** Correlation of human H3F3A and H3F3B codon usage frequencies with those computed for the genes associated with cell proliferation (orange) and cell differentiation (green) [41]. Each dot represents individual gene from the corresponding group. The dotted lines indicate the correlation coefficient medians for each group and the H3.3 gene. **C.** Benchmarking of the codon usage frequencies in the H3.3 genes relative to the frequencies estimated for the genes from the cell proliferation and differentiation groups. Boxplots represent correlation values for the amino-acid specific codon frequencies of the individual cell proliferation genes or cell differentiation genes with the overall profiles estimated for their own groups. Dashed lines show the mean values of the correlations of individual genes from one group with the opposite group profile (e.g. mean for the correlations of the codon usages of the proliferation genes with the overall differentiation profile). The brown and blue diamonds indicate correlation values for the human H3F3A and H3F3B genes. **D.** A model illustrating the possible role of the evolutionary conserved arrangement of the two genes (H3F3A and H3F3B) encoding the same protein (H3.3) in fine-tuning of this protein expression.

To gain further insight on the evolutionary adaptation of the H3.3 genes, we compared their codon usage frequencies to those estimated for the two groups of genes shown to be involved in different transcriptional programs (‘cell proliferation’ and ‘cell differentiation’ genes; data from [41]). Specifically, we computed pairwise correlation between the amino-acid specific codon frequencies of the H3.3 genes and the individual genes associated with each of transcriptional program (orange and green dots in Figs. 5B, S7B). This analysis showed that, by this metric, *H3F3A* shares greater similarity with the ‘proliferation’ genes, while *H3F3B* is more similar to the ‘differentiation’ genes (P = 6.91*10^−12^ and P = 8.3*10^−12^ respectively, Mann-Whitney’s test; Fig S7C-D). As previously, we confirmed these results in a similar analysis based on direct codon frequencies which are not corrected for amino-acid abundance (Fig. S7E-F).

To benchmark the similarity between the codon usage of an individual gene and the codon usage profiles associated with different transcriptional programs, we correlated codon usages of individual proliferation‐ and differentiation-induced genes to both codon usage profiles (Fig. 5C). Comparison of the H3.3 genes with these benchmarks showed that *H3F3A* falls within 25^th^ percentile of the proliferation-associated genes when they are evaluated against codon usage profile of their own group (r=0.58). The similarity of this gene to the differentiation group is low and it is on par with the average similarity observed for the proliferation-induced genes when they are compared to the codon usage profile of the differentiation group. In line with our previous results, *H3F3B* exhibits an opposite trend: its codon usage correlates better with differentiation gene profile (r=0.71 vs. r=0.35 for differentiation and proliferation profiles respectively). We note however, that the *H3F3B* ranks relatively lowly among the differentiation-induced in terms of their similarity to the group profile.

Based on these results, we conclude that *H3F3A* and *H3F3B* were evolutionary optimized for distinct transcriptional programs. In this analysis we tested two programs that have been described in literature [41]. While other programs may exist, our observations indicate better fitness of *H3F3A* for the proliferation program and, arguably to a lesser extent, better fitness of *H3F3B* for differentiation program. We also found that, similar to *H3F3B* (but not *H3F3A*), the differentiation-induced genes correlate strongly with the genome-wide codon usage (r=0.88), which suggests a broad expression profile. Thus, while *H3F3B* does not rank high within the differentiation-induced genes, taken together our findings show that this gene is broadly expressed in cell types, including differentiated cells. Overall, we report that despite encoding identical protein sequence, *H3F3A* and *H3F3B* have distinct evolutionary histories and are optimized for distinct transcriptional programs at the codon usage level, as illustrated in Figure 5D.

## Discussion

The H3.3 histone is currently a subject of intense research due to its biological and biomedical significance [21,22]; however, evolution of the genes encoding this protein is not fully understood. In this study, we addressed this issue and studied the evolutionary history of the H3.3-encoding genes from a diverse set of metazoan genomes. All analyzed genomes harbor multiple genes (two in most cases, *H3F3A* and *H3F3B*) that encode an identical protein sequence. We have shown that, despite being highly conserved at the protein level, H3.3-encoding genes are subject to selection pressure at DNA sequence level, which is related to their cellular function. Several lines of evidence stemming from phylogenetic analysis, as well as analyses of the gene structure, synteny and codon usage (Figs 1, 2, 3 and 5) indicate that *H3F3A* is specific for sarcopterygian (lobe-finned fish) lineage, whereas *H3F3B* exist in all sarcopterygians and bears similarity to H3.3 genes in actinopterygians (ray-finned fish) and jawless fish and with the vertebrate RD H3.1/H3.2 genes that diverged much earlier. These results suggest that *H3F3B* is more similar to ancestral form of H3.3 gene than *H3F3A*, which could be a product of a duplication event occurring after actinopterygian-sarcopterygian split. However, we cannot completely exclude that *H3F3A* could have been lost in actinopterygians and other lineages and additional studies are required to exactly trace the origin of each H3.3 gene.

Despite absolute conservation at the protein level in both genes, tetrapod *H3F3A* and *H3F3B* are under varying degrees of purifying selection at the codon synonymous sites, resulting in distinct codon usage profiles in these genes (Fig. 5). Our analysis revealed that codon usage in *H3F3B* is similar that of the ‘cell differentiation-induced’ genes, in contrast to the codon usage in *H3F3A*, which is similar to that of ‘cell proliferation-induced genes’ [41]. We note that while proliferation-induced genes are active in a specific pathway, one can expect that the ‘differentiation-induced’ genes would show a broad expression profile as a group, because they can be associated with various pathways in different cell types. This is also in line with our observation that codon usage of *H3F3B*, but not of *H3F3A*, is similar to that of UEGs which are active throughout cell types (Fig. 5B). Furthermore, similarly to the UEGs, *H3F3B* genes feature a compact structure, with short introns (Fig. 1B) [45,46]. Given that we analyzed only two transcriptional programs, it is possible that *H3F3A* and/or *H3F3B* would show similar or even better fit for other programs. However, our results allow us to conclude that *H3F3A* and *H3F3B* genes are evolutionary optimized for different transcriptional programs through codon usage preferences and intron-exon organization.

In summary, the H3.3 genes provide a unique ‘study case’, in which protein sequence remain constant in course of evolution for an extended time period, allowing analysis of the selection operating at nucleotide level. Such analysis reveals an evolutionary mechanism of nucleotide sequence optimization for fine-tuning of gene expression in specific cellular programs.

## Methods

### Phylogenetics analysis

Sequences and annotations of the genes encoding histone variant H3.3 in different species, as well as other genes used in this study were obtained from Ensembl and NCBI-RefSeq databases. A phylogenetic tree was constructed using PHYML3.1 software [47], with approximate likelihood ratio test (Chi2-based) for branch supports and GTR nucleotide substitution model.

### Synteny analysis

Synteny around *H3F3A* and *H3F3B* genes in selected set of vertebrate genomes was detected using a web application Genomicus version 80.01, that uses Ensembl comparative genomic data (http://genomicus.biologie.ens.fr/genomicus) [32]. To supplement Genomicus-based analysis and test for synteny between tetrapods and distant organisms such as fly and lamprey, an additional method was used. This method measures the degree of conservation of the genes neighboring H3.3-encoding genes based on comparison of their CDS and translated amino-acid sequences. The annotated chromosome sequences were downloaded from Ensembl (http://www.ensembl.org/info/data/ftp/index.html). Biopython (www.biopython.org) was used to extract CDS of 30 genes upstream and downstream of all tetrapod *H3F3A* and *H3F3B*, and of the H3.3 genes in distant organisms: actinopterygians (tetraodon, zebrafish and medaka), lamprey and fly. Pairwise comparison of nucleotide and protein sequences was done by aligning two sequences using MUSCLE [48] and computing sequence identity scores.

### 3’UTRs comparison

3’UTR sequences of actinopterygian H3.3 genes were compared to those of tetrapod *H3F3A* and *H3F3B* to find similarities. Comparison was performed through alignment of each pair of 3’UTR sequences using MUSCLE and computing their sequences identity scores with gap exclusion. Since gaps (indels) in alignments can substantially influence final identity scores [49], we excluded them from calculations to insure that high UTR sequence variability due to insertions and deletions does not deflate the scores and affect comparisons.

### Codon usage analysis

Two metrics of codon usage were used, the ‘amino-acid specific codon frequencies’ and ‘codon frequencies’. The amino-acid specific codon frequencies represent codon occurrences normalized for amino-acids abundance [41], i.e. divided by the number of times the corresponding amino-acid appears in the protein sequence. This metric corrects for potential amino-acid usage biases and represents the probability that a codon will be used given that the corresponding amino-acid is used. The second metric, ‘codon frequencies’, were computed by dividing the codon occurrences by the total number of codons in the gene (i.e. normalized by the length of the encoded amino-acid sequence). The codon usage profiles were computed for different gene sets (proliferation-induced [41], differentiation-induced [41]). Genome-wide codon counts were obtained from (http://www.kazusa.or.jp/codon).

## Acknowledgements

We thank Marjorie Oettinger for valuable discussions and Mattia Lion, Behfar Aldehali, and Erica Larschan for critical reading of the manuscript and many insightful comments.

